# Receptor Elimination by E3 Ubiquitin Ligase Recruitment (REULR): A Targeted Protein Degradation Toolbox

**DOI:** 10.1101/2022.10.31.514624

**Authors:** Dirk H. Siepe, Lora K. Picton, K. Christopher Garcia

## Abstract

In recent years, targeted protein degradation (TPD) of plasma membrane proteins by hijacking the ubiquitin proteasome system (UPS) or the lysosomal pathway have emerged as novel therapeutic avenues in drug development to address and inhibit canonically difficult targets. While TPD strategies have been successful to target cell surface receptors, these approaches are limited by the availability of suitable binders to generate heterobifunctional molecules. Here, we present the development of a nanobody (VHH) based degradation toolbox termed REULR (Receptor Elimination by E3 Ubiquitin Ligase Recruitment). We generated human and mouse cross-reactive nanobodies against 5 transmembrane PA-TM-RING type E3 Ubiquitin Ligases (RNF218, RNF130, RNF167, RNF43, ZNRF3) covering a broad range and selectivity of tissue expression with which we characterized expression in human and mouse cell lines and immune cells (PBMCs). We demonstrate that heterobifunctional REULR molecules can enforce transmembrane E3 ligase interaction with a variety of disease relevant target receptors (EGFR, EPOR, PD-1) by induced proximity resulting in effective membrane clearance of the target receptor at varying levels. In addition, we designed E3 Ligase self-degrading molecules, ‘fratricide’ REULRs (RNF128, RNF130, RENF167, RNF43, ZNRF3), that allow downregulation of one or several E3 Ligases from the cell surface and consequently modulate receptor signaling strength. REULR molecules represent a VHH-based, modular and versatile “mix and match” targeting strategy for facile modulation of cell surface proteins by induced proximity to transmembrane PA-TM-RING E3 ligases.

## MAIN

Classical drug discovery approaches against membrane protein targets such as cell surface receptors generally rely on small molecule inhibitors and monoclonal antibodies, a vast majority of disease relevant cell surface receptors still remains extremely challenging to target and have been largely deemed ‘undruggable’ by established screening strategies (Mullard, 2019). Finding alternative strategies to target challenging plasma membrane proteins has therefore become a prime focus in recent years.

Targeted protein degradation has emerged as a novel therapeutic strategy in drug development by directing proteins to the cells own degradation machinery (UPS) (Békés et al., 2022; Deshaies, 2015; Sakamoto et al., 2001). The majority of degraders such as PROTACs (Sakamoto et al., 2001), Molecular Glues (Dong et al., 2021), dTags (Nabet et al., 2018), or TRIM-Away (Clift et al., 2017) are based on a heterobifunctional design that leads to the formation of a ternary complex between a cytosolic E3 ubiquitin Ligase and a protein of interest to facilitate ubiquitination and subsequent 26S proteasome dependent degradation (Li and Crews, 2022). While classical degraders have been successful (Mullard, 2019), this approach is ultimately limited to cytosolic targets and therefore 1/3 of the protein-coding genes representing the membrane proteome is not accessible by this approach (Thul et al., 2017; Uhlen et al., 2015).

More recently, targeted protein degradation approaches utilizing lysosomal degradation strategies (LYTAC, KineTac) (Banik et al., 2020; Pance et al., 2022) and proteolysis-targeting antibodies (AbTac, ProTab) using WNT related transmembrane E3 ligases (RNF43, ZNRF3) have emerged (Cotton et al., 2021; Marei et al., 2022). These approaches tether target proteins on the cell surface to either lysosome shuttling receptors or cell-surface E3 ubiquitin ligases to induce membrane clearance. Both technologies are mainly limited by the availability and specificity of shuttling receptors or transmembrane E3 binding moieties, selectivity (tissue expression), and design (antibody formatting) and complexity of production.

In an effort to accelerate the development of targeted protein degradation tools, we present a modular and versatile nanobody (VHH) based protein degradation toolbox termed REULR (Receptor Elimination by E3 Ubiquitin Ligase Recruitment). We generated human and mouse cross-reactive nanobodies against ECDs (extracellular domain) of 5 transmembrane PA-TM-RING type E3 Ubiquitin Ligases (RNF218, RNF130, RNF167, RNF43, ZNRF3) covering a broad range and selectivity of tissue expression. Next, we utilized our VHHs to characterize expression of these 5 PA-TM-RING E3 Ligases in commonly used human and mouse cell lines and immune cells (T cells, Monocytes, B cells, NK cells). We demonstrate that heterobifunctional REULR molecules can enforce transmembrane E3 ligase interaction with a variety of disease relevant target receptors (EGFR, EPOR, PD-1) by induced proximity resulting in robust membrane attenuation of the target receptor. Furthermore, we present a strategic approach to tune transmembrane E3 Ligases itself by generating homo-, heterobifunctional and arrayed multimeric Fratricide REULRs and consequently modulate signaling events of natural target receptors.

### PA-TM-RING E3 Ligase Nanobodies for Receptor Elimination

The human transmembrane (TM) E3 ligase family represents a class of diverse RING type E3 ubiquitin ligases (Cai et al., 2022; Deshaies and Joazeiro, 2009) with approximately 50 members (Fig.1A upper chart). These proteins exert widespread involvement in several diseases and cancer (Cai et al., 2022; Chen et al., 2022). The family can be further grouped into subcellular and structurally related sub classes, the plasma membrane localized E3 TM ligases include: RING domain containing proteins (7), PA-TM-RING (10), RING between RING (RBR; 5) and the membrane-associated RING-CH (MARCH; 4) families (Figure 1A, lower chart). In general, E3 Ligases are notoriously challenging to study and their substrates still remain highly elusive, mostly due to the nature of the ubiquitylation cascade, which are characterized by very weak target affinities and fast kinetics (Duan and Pagano, 2021; Komander and Rape, 2012; Metzger et al., 2014).

**Figure 1.**
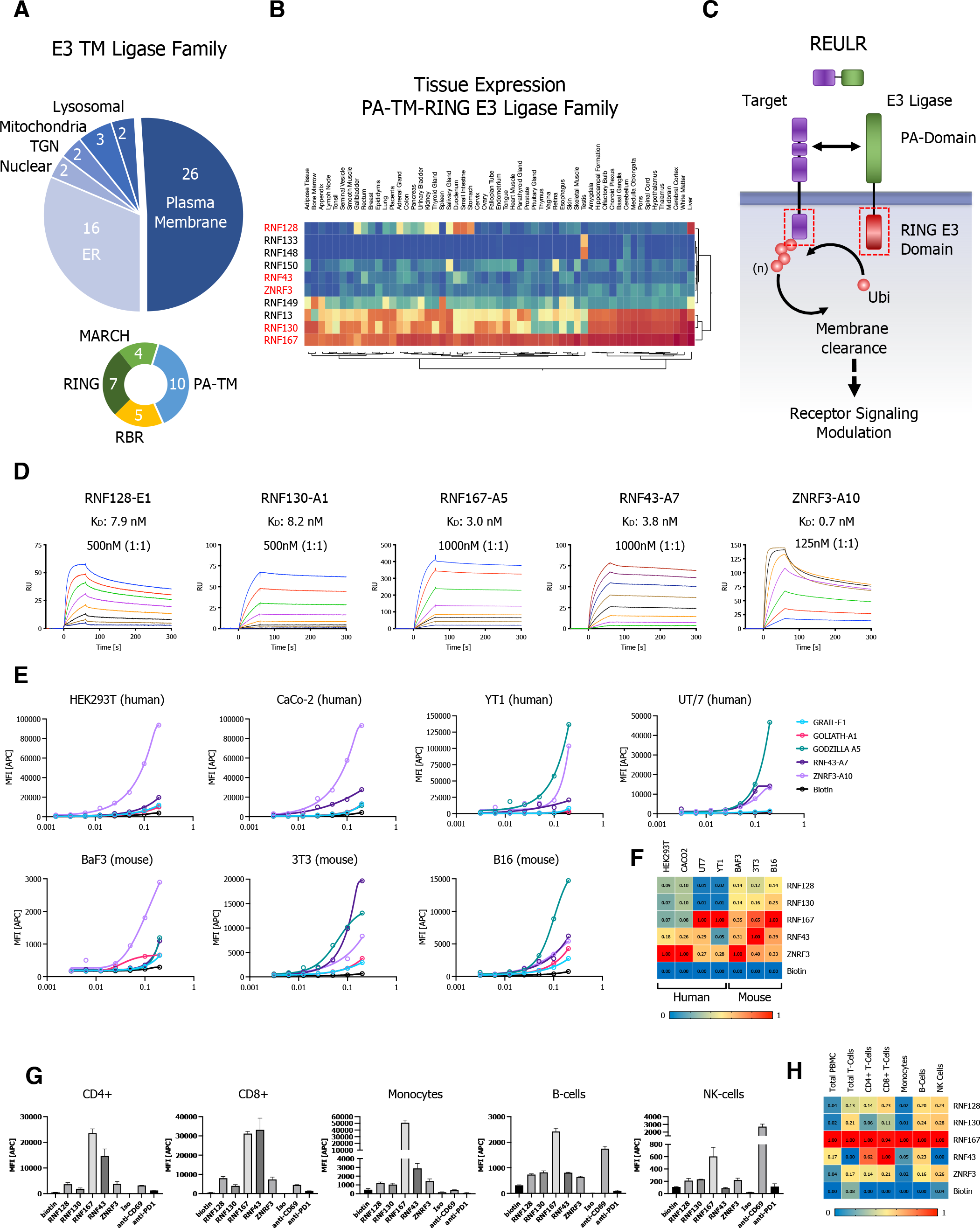
Transmembrane PA-TM-RING E3 Ligase nanobodies for Receptor Elimination. (A) Pie chart representation of the transmembrane E3 Ligase family classified by subcellular localization (upper chart). Plasma membrane localized transmembrane E3 Ligase subfamily, grouped into subcellular and structurally related sub classes (lower char). (B) Hierarchical 2-way clustering heatmap of normal tissue mRNA expression data for the PA-TM-RING E3 Ligase subfamily. (C) Schematic representation of the REULR concept. Enforced transmembrane E3 ubiquitin Ligase recruitment to a target receptor reduces target receptor cell surface levels by E3 Ligase dependent intracellular ubiquitination and subsequent membrane clearance. (D) SPR sensograms and binding affinities of PA-TM-RING ligase selected nanobodies (analytes) for human RNF128, RNF130, RNF167, RNF43 and ZNRF3 ECDs (ligands). (E) Cell surface staining of representative human (HEK239T, CaCo-2, YT1, UT/7) and mouse (BaF3, 3T3, B16) cell lines using a panel of 5 PA-TM-RING E3 Ligase binding nanobodies (Nanobody:SA647 tetramers) and analysis by flow cytometry, full titration (1:1 dilutions; 200nM tetramer), Biotin:SA647 (Biotin) served as a negative control. (F) Staining data visualized in a normalized heatmap for human and mouse cell lines. (G) PBMC (Peripheral blood mononuclear cells) immunophenotyping panel to identify binding of 5 PA-TM-RING E3 Ligase binding nanobodies (Nanobody:SA647 tetramers; 200nM) to T cells (CD4+; CD8+), Monocytes, B cells and NK cells, analysis by flow cytometry. Biotin:SA647 served as a negative control (Biotin). Anti-PD1, anti-CD69 were used as phenotyping control antibodies in comparison to an Isotype control. (H) PBMC sub cell type data summarized in a normalized heatmap. Data are represented as mean ± SD (n=3).

Here, we focused on the PA-TM-RING type E3 ligases (Nakamura, 2011), a family of approximately 10 members with a broad tissue expression pattern (Figure 1B) and a unique domain architecture that is minimally defined by three conserved domains: A extracellular protease-associated (PA) domain that acts as a substrate recruitment domain, a transmembrane domain (TM), and a cytosolic catalytic RING type E3 ligase domain (RING-H2 finger; RNF) (Figure 1C). Mechanistically, the cytosolic RING E3 ligase domain functions as an allosteric activator and scaffold that recruits the ubiquitin machinery in close proximity to a substrate while the extracellular PA domain functions as a substrate recruitment domain. We therefore hypothesized that PA-TM-RING E3 ligases could be retasked to selectively eliminate non-natural cell surface targets by an induced proximity approach that we termed REULR: Receptor Elimination by E3 Ubiquitin Ligase Recruitment (Figure 1C).

To develop a modular and versatile toolbox, we first identified 5 PA-TM-RING E3 ligases covering a wide range of tissue expression allowing cell type specific REULR approaches: RNF128 (GRAIL), RNF130 (GOLIATH), RNF167 (GODZILLA), RNF43 and ZNRF3 (Figure 1B; marked in red). Therapeutic monoclonal antibodies (mABs) and antibody engineering have revolutionized cancer therapies in the last decade (Robert, 2020) but they are not without limitations, mainly size, complexity of formatting, expression and modularity. In order to overcome these limitations, we took advantage of the superior pharmacokinetic properties of nanobodies (VHH) such as small size (1/10 the size of conventional antibodies), high stability, strong antigen-binding affinity, modularity and ease of expression (Bannas et al., 2017; Jin et al., 2022; Yang and Shah, 2020). We screened a synthetic nanobody library allowing rapid high-throughput selection by yeast display (McMahon et al., 2018) using the ECDs (extracellular domains) of human RNF128 (GRAIL), RNF130 (GOLIATH), RNF167 (GODZILLA), RNF43 and ZNRF3 that led to 8 Nanobodies against 5 Ligases with nanomolar to picomolar affinities (Figure 1D, Figure 1-figure supplement 1A-C).

A pairwise protein sequence alignments of the human and mouse ECDs of the 5 PA-TM-RING type E3 Ligases revealed that the ECDs are highly conserved between both species, with an average amino acid sequence identity of 97.75% (Figure 1-figure supplement 1D). We therefore tested our PA-TM-RING E3 Ligase nanobodies against a panel of commonly used human (HEK293T, CaCo-2, YT1, UT/7) and mouse (BaF3, 3T3, B16) cell lines by cell surface staining as indicated (Figure 1E, Figure 1-figure supplement 2A, B) and summarized in a normalized heatmap (Figure 1F). Indeed, all nanobodies tested were cross reactive against human and mouse cell lines which poses a significant advantage for the design and application of REULR molecule for in vitro and in vivo studies. We next evaluated the nanobodies on primary cells using PBMCs (Primary Peripheral Blood Mononuclear Cells) to identify cell surface binding to immune cells: T cells (CD4+; CD8+), Monocytes, B cells and NK cells (Figure 1G, Figure 1-figure supplement 2C), summarized in normalized heatmap (Figure 1H). Similar to human and mouse cell lines, we observed some ligases like RNF167 being highly expressed in many cell types while most other ligases tested show a more nuanced, cell type specific expression pattern (Figure 1F, H).

### Receptor Elimination by E3 Ubiquitin Ligase Recruitment (REULR)

To evaluate potential targets for our REULR approach we performed a membrane proteome wide analysis of reported ubiquitin sites (Hornbeck et al., 2015). On average, 45% of cell surface receptors were reported to have at least one or more ubiquitin site (Figure 2A) which represents an untapped potential for cell surface receptors to be targeted using a REULR strategy. Members of the receptor tyrosine kinases (RTK) family including cytokine receptors EpoR (via JAK2 *V617F*) and members of the epidermal growth factor receptor (ErbB; HER) family represent the most common oncogenic drivers of malign carcinomas (Gschwind et al., 2004; Lemmon and Schlessinger, 2010; Schmidt-Arras and Böhmer, 2020). However, despite their immense clinical relevance, conventional drug discovery approaches have shown limited efficacy and problems of resistance (Chong and Jänne, 2013; Yang et al., 2022). These limitations are mainly due to the nature of primary and emerging secondary escape mutations in the receptor (EGFR *T790M*) and acquired resistance as well as pathway mutations e.g. JAK2 *V617F* (EPOR/TPOR) that lead to constitutively over activation and dysregulation with detrimental outcome for patients (Chan et al., 2017; Glassman et al., 2022; Kumar et al., 2020; Shi et al., 2022).

**Figure 2.**
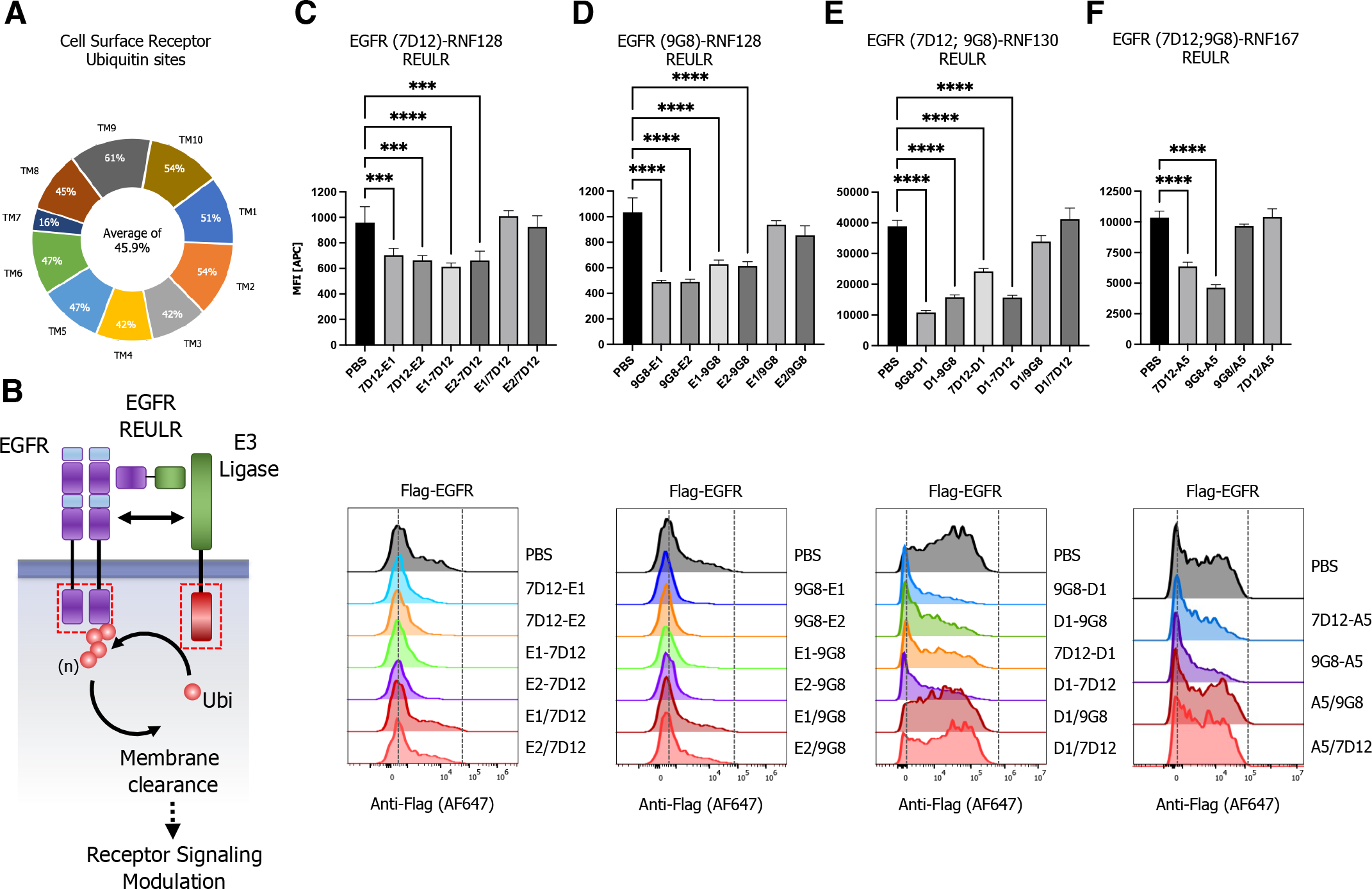
Epidermal Growth Factor Receptor (EGFR) Elimination by E3 Ligase Recruitment (REULR). (A) Analysis of MS (Mass Spectrometry) validated proteome wide ubiquitin sites matched to the human membrane proteome, subclassified by number of transmembrane domains. (B) Schematic representation of EGFR degradation using a EGFR-REULR molecule. (C-F) HEK293T cells were transiently transfected with FLAG-tagged full-length EGFR cDNA (human) under the control of a constitutively active CMV (cytomegalovirus) promoter. 24h post transfection, cells were incubated with EGFR-REULR molecules (50nM) as indicated using RNF128 (E1; E2), RNF130 (D1) or RNF167 (A5) targeting nanobodies in combination with EGFR binding moieties (7D12; 9G8) in varying orientations as indicated in comparison to monomeric nanobodies or PBS. After 24h cells were subjected to FACS analysis using a FLAG antibody (Alexa Fluor 647 Conjugate) to monitor EGFR levels on the cell surface. Representative FACS histograms are visualized below the quantified data. Data are mean ± s.d. (n = 3 replicates).

We first designed different combinations of heterobifunctional REULRs to EGFR using two VHH (7D12; 9G8) that were previously described to inhibit ligand binding to EGFR: Nanobody 7D12 sterically blocks ligand binding to EGFR similar to cetuximab and 9G8 acts by inhibiting high affinity ligand binding and dimerization (Roovers et al., 2011; Schmitz et al., 2013; Wolf et al., 2007). To assess whether EGFR can be degraded by this set of REULR molecules, we overexpressed FLAG-tagged full-length EGFR in HEK293T cells that endogenously express PA-TM-RING E3 ligases at varying levels (Figure 1E) and treated cells with intact REULR molecules, monomeric versions or PBS served as negative controls (Figure 2C-F, Figure 2-figure supplement 1A-D). We observed EGFR degradation after treatment with EGFR-REULR molecules at varying efficiency depending on choice of E3 targeting ligase, EGFR VHH and orientation (Figure 2C-F). Collectively, EGFR-REULR designs using the N-terminal 9G8 nanobody in combination with C-terminal RNF128, RNF130 or RNF167 targeting nanobodies performed better and resulted in more effective EGFR degradation compared to other designs.

To show modularity with other binding moieties, we reformatted an EpoR targeting Diabody (DA10) (Moraga et al., 2015) into an scFv (single-chain variable fragment) and fused it to RNF128, RNF43 and ZNRF3 targeting nanobodies (Figure 3A). Indeed, intact EPOR-REULR molecules could efficiently degrade EPOR while showing no activity when cells were treated with monomeric version of the individual targeting arms or PBS (Figure 3B-D, Figure 3-figure supplement 1A). Of note, degradation efficiency did not directly correlate with expression levels of PA-TM-RING E3 ligases observed in HEK293T cells. While RNF128 and RNF43 appear to be expressed at much lower levels than ZNRF3 (~25x; Figure 1E), degradation using EPOR-RNF128 or EPOR-RNF43 REULR (Figure 3B, C) still resulted comparable levels of EPOR loss in comparison to a ZNRF3 based EPOR-REULR molecule (Figure 3D).

**Figure 3.**
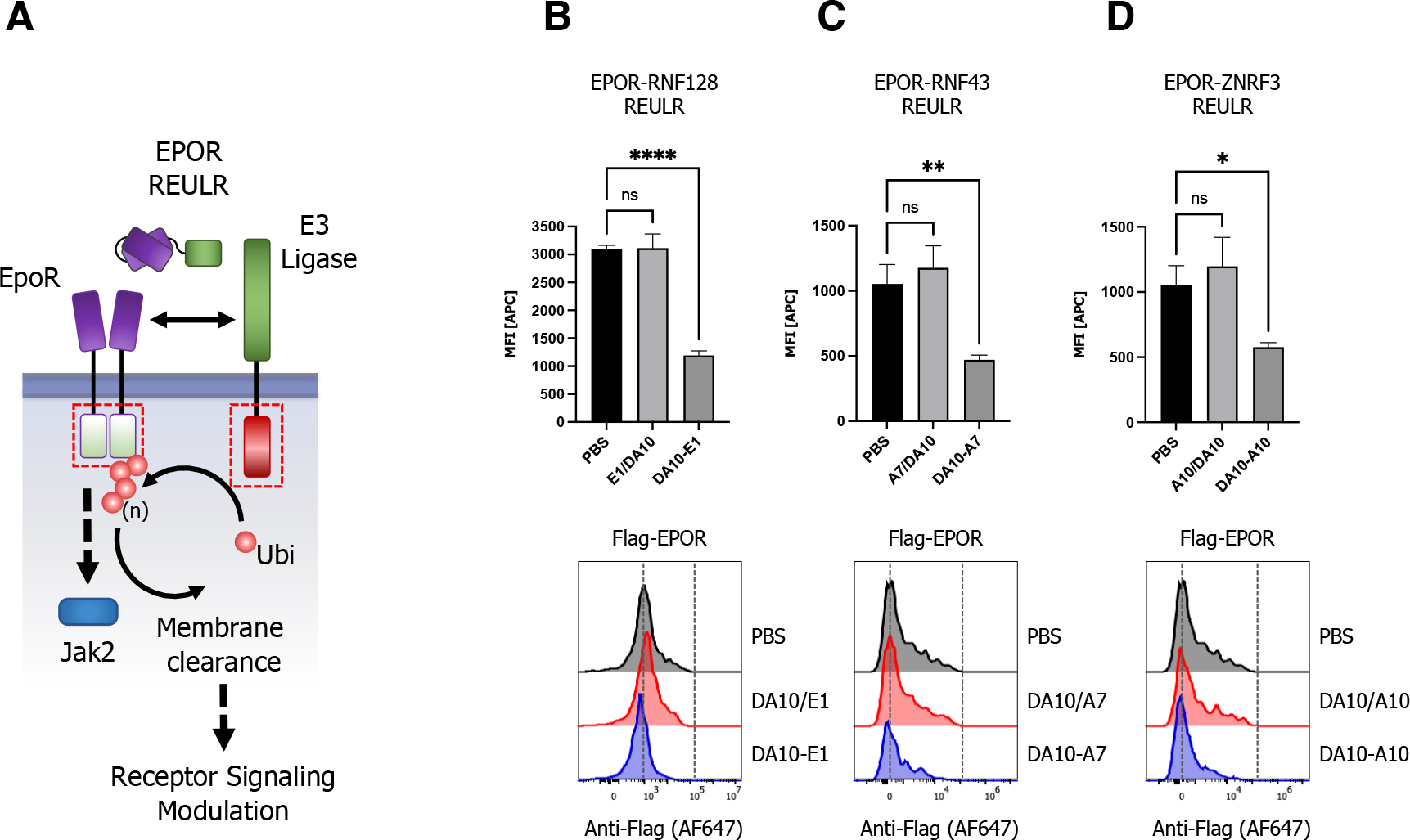
EpoR Elimination by E3 Ligase Recruitment (REULR). (A) Schematic representation of EPOR-REULR mediated EPOR degradation. (B-D) HEK293T cells were transiently transfected with FLAG-tagged full-length EPOR cDNA (human) under the control of a constitutively active CMV (cytomegalovirus) promoter. 24h post transfection, cells were incubated with EPOR-REULR molecules (50nM) as indicated using RNF128 (E1), RNF43 (A7) or ZNRF3 (A10) targeting nanobodies fused to a scFv (Single Chain Fragment Variable) reformatted EPOR Diabody (DA10). Monomeric binding moieties or PBS were used as negative control. After 24h cells were subjected to FACS analysis using a FLAG antibody (Alexa Fluor 647 Conjugate) to monitor EPOR levels on the cell surface. Representative FACS histograms are visualized below the quantified data. Data are mean ± s.d. (n = 3 replicates).

Immunotherapies based on checkpoint biology have emerged as a major pillar in fighting cancer. Immune-checkpoint inhibitors (ICIs) such as antibodies targeting CTLA-4 (ipilimumab), PDL1 (atezolizumab and durvalumab) or PD1 (pembrolizumab and nivolumab) have become some of the most widely used anticancer therapies (Johnson et al., 2022; Robert, 2020). However, immune-related adverse events (irAEs), such as autoimmune symptoms and tumor hyperprogression, present a significant challenge in the clinic (Conroy and Naidoo, 2022) and a need for a continuous development of immune-oncology pipeline drugs. Targeted protein degradation could provide a major expansion in the repertoire of modulating immune checkpoint receptor by directly regulating cell surface levels. We therefore next generated REULR molecules targeting the immune checkpoint receptor PD-1 (Programmed cell death protein 1) by fusing an anti-human PD-1 nanobody (Fernandes et al., 2020) to several nanobodies targeting RNF128, RNF130 and RNF167 (Figure 4A-D). Similar to EGFR- and EPOR-REULRs, treatment of HEK293T cells overexpressing FLAG-tagged full-length PD-1 with a variety of PD1-REULR molecules using RNF128, RNF130 or RNF167 targeting nanobodies resulted in robust and near complete loss of PD-1 receptor from the cell surface compared to treatment with the respective monomeric VHHs or PBS (Figure 4B-D, Figure 4-figure supplement 1A). While RNF130 based REULR molecules worked most effectively in degrading EGFR, PD1-REULR molecules using RNF128 and ENF167 targeting nanobodies collectively resulted in near-complete elimination of PD1.

**Figure 4.**
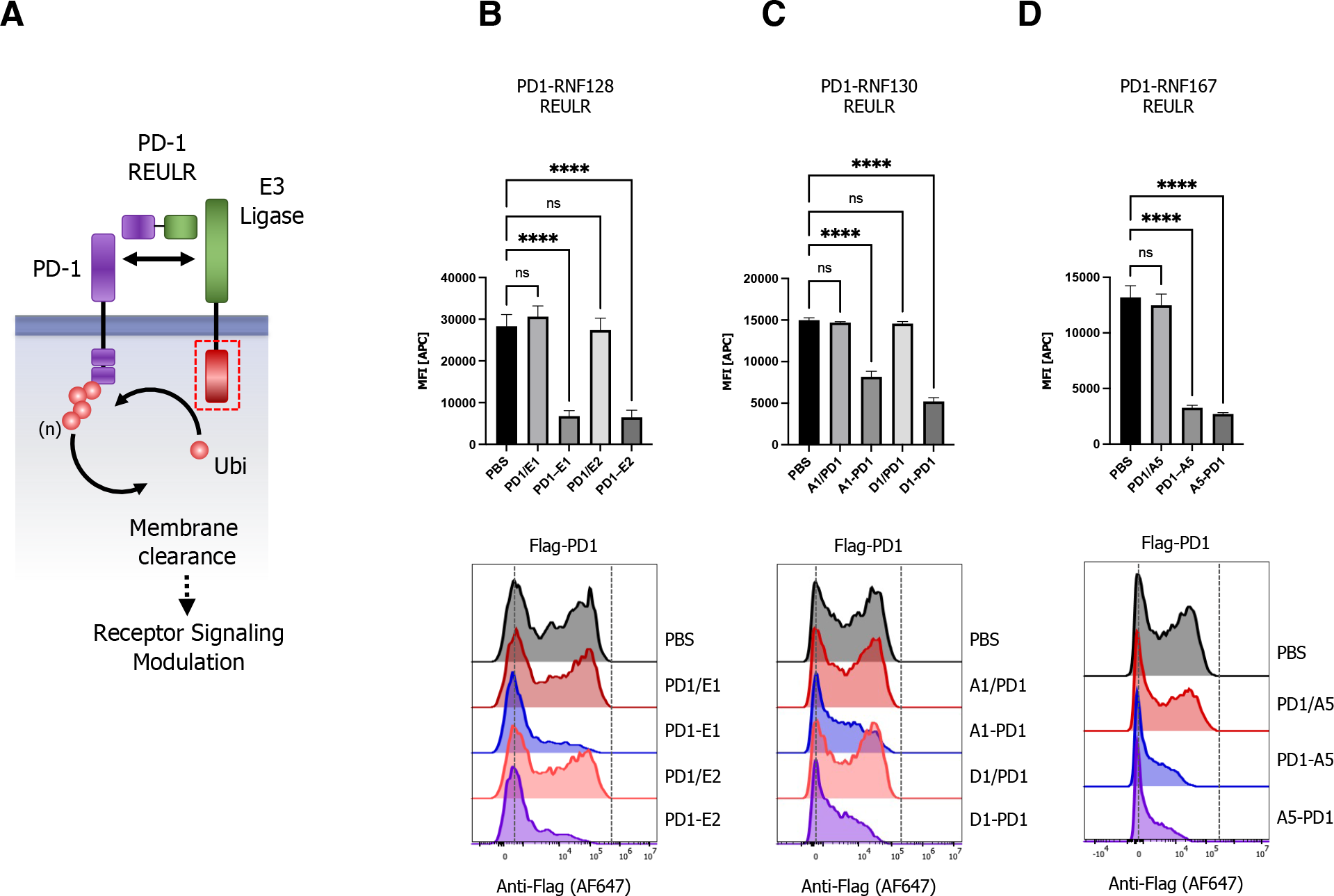
Immune Checkpoint Receptor PD-1 Elimination by E3 Ligase Recruitment (REULR). (A) Schematic representation of PD1-REULR mediated PD-1 degradation. (B-D) HEK293T cells were transiently transfected with FLAG-tagged full-length PD-1 cDNA (human) under the control of a constitutively active CMV (cytomegalovirus) promoter. 24h post transfection, cells were incubated with PD1-REULR molecules (50nM) as indicated using RNF128 (E1; E2), RNF130 (A1; D1) or RNF167 (A5) targeting nanobodies fused to a PD-1 binding nanobody (PD1). Monomeric binding moieties or PBS were used as negative controls. After 24h cells were subjected to FACS analysis using a FLAG antibody (Alexa Fluor 647 Conjugate) to monitor PD-1 levels on the cell surface. Representative FACS histograms are visualized below the quantified data. Data are mean ± s.d. (n = 3 replicates).

### Expansion of the REULR platform to modulate E3 Ligases itself: Fratricide REULRs

Emerging evidence highlights the pivotal role of RING-type E3 ligases and their substrates in a wide range of human diseases and mutation of RING-type E3s, or modulation of their activity is frequently associated with pathogenesis including viral infections, neurodegenerative disorders, autoimmune diseases and cancer (Cai et al., 2022; Chen et al., 2022; Duan and Pagano, 2021). Indeed, RNF43 mutations have been associated with aggressive tumor biology such as colorectal and endometrial cancer (Fang et al., 2022; Matsumoto et al., 2020; van Herwaarden et al., 2021). To evaluate the impact of other PA-TM-RING E3 ligases in cancer we analyzed TCGA (The Cancer Genome Atlas) tissue mRNA expression data obtained for RNF128, RNF130, RNF167, RNF43 and ZNRF3 from 17 cancer types representing 21 cancer subtypes. The data shows elevated expression of E3 ligases in various cancer types. Notably, while RNF167 is highly expressed in almost all cancers, other ligases like RNF128 (thyroid, liver, urothelial, colorectal), RNF130 (glioma) or RNF43 (colorectal cancers) show a more selective tissue associated expression pattern (Figure 5A) in cancer cells.

**Figure 5.**
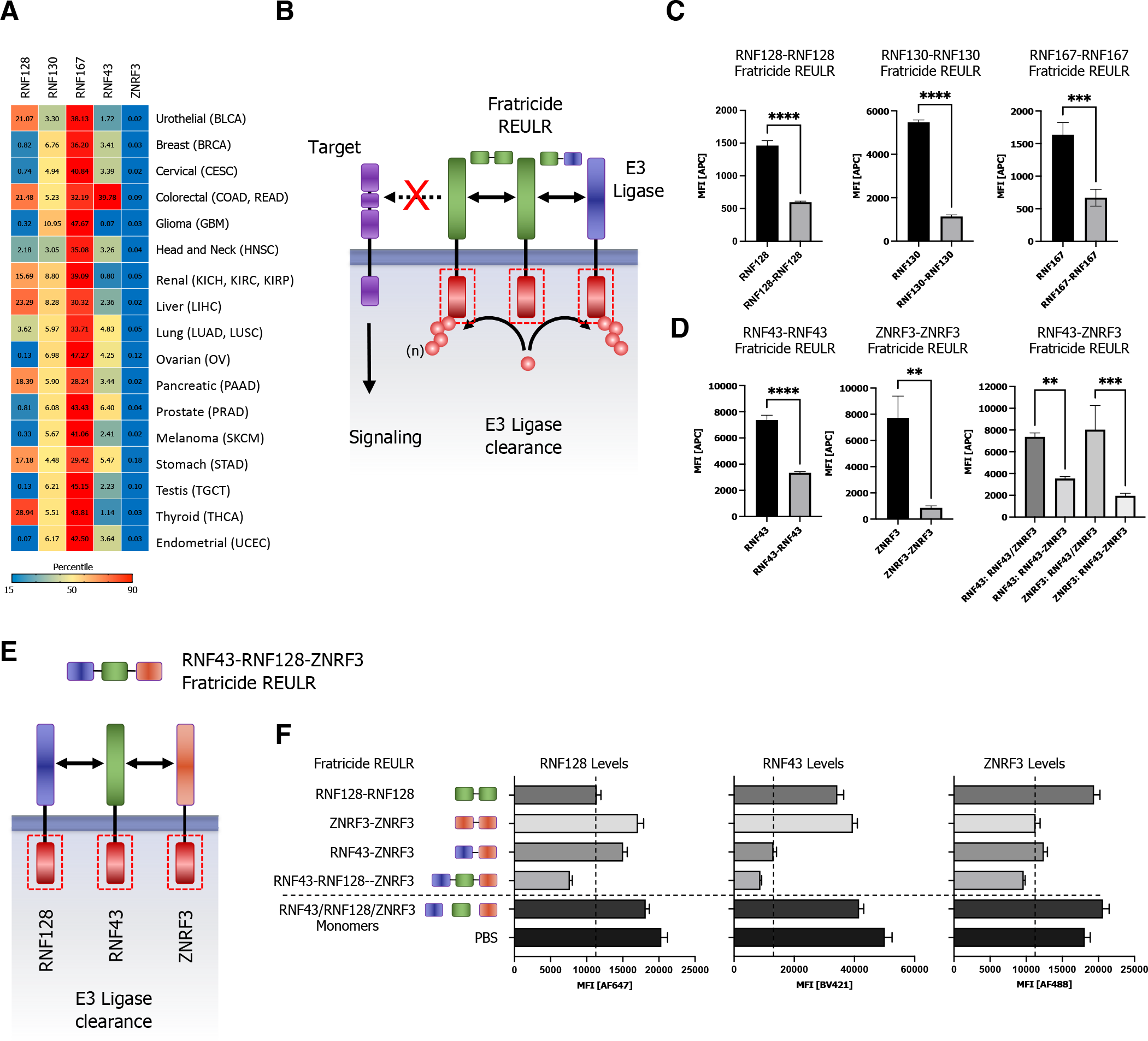
Homo- and heterobifunctional Fratricide REULR. (A) TCGA cancer tissue RNA-seq data for RNF128, RNF130, RNF167, RNF43 and ZNRF3 was obtained for all 20 screened secreted ligands from 17 cancer types representing 21 cancer subtypes and were processed as median FPKM (number Fragments Per Kilobase of exon per Million reads) and visualized as a hierarchical clustering heatmap. (B) Schematic representation of homo or heterobispecific Fratricide REULR. (C) HEK293T cells were transiently transfected with HA-tagged full-length PA-TM-RING E3 Ligase cDNA (human) under the control of a constitutively active CMV (cytomegalovirus) promoter, as indicated. 24h post transfection, cells were incubated with Fratricide REULR molecules (50nM) targeting RNF128, RNF130, RNF167, RNF43 or ZNRF3. (D) HEK293T cells were co-transfected with HA-tagged full-length RNF43 and MYC-tagged full-length ZNRF3 cDNA (human) and treated with a heterobispecific RNF43-ZNRF3 Fratricide REULR 24h post transfection (monomeric binding moieties were used as negative controls). After 24h cells were subjected to FACS analysis using a HA antibody (Alexa Fluor 647 Conjugate) or MYC antibody (Alexa Fluor 488 Conjugate) to monitor PA-TM-RING E3 Ligase levels on the cell surface. Data are mean ± s.d. (n = 3 replicates). (E) Schematic representation of heterobispecific arrayed multimeric Fratricide REULR. (F) HEK293T cells were co-transfected with HA-tagged full-length RNF128, MYC-tagged full-length ZNRF3 and FLAG-tagged full-length RNF43 cDNA (human). 24h post transfection cells were treated with homo- or heterobispecific Fratricide REULR molecules as indicated, or a RNF43-RNF128-ZNRF3 multimeric Fratricide REULR (PBS and monomeric binding moieties were used as negative controls). After 36h cells were subjected to FACS analysis using a HA antibody (Alexa Fluor 647 Conjugate), MYC antibody (Alexa Fluor 488 Conjugate) and FLAG antibody (Brilliant Violet 421) to monitor RNF128 (left panel), RNF43 (middle) and ZNRF3 (right panel) E3 Ligase levels on the cell surface. Data are mean ± s.d. (n = 3 replicates).

Despite their critical role in regulating protein homeostasis and pathological signaling, our understanding of transmembrane E3 ligase mediated signaling still remains largely fragmented and can mainly be attributed to the limited availability of tools to study TM E3 ligases. Interestingly, the activity of E3 ligases is tightly regulated by post-translational modifications and a typical feature of most ligases is the ability to catalyze their own ubiquitination (de Bie and Ciechanover, 2011; Lorick et al., 1999). Based on this paradigm, we postulated that self-regulation by auto-ubiquitination could be used to regulate E3 ligase dependent signaling.

We therefore proceeded in developing REULR molecules that target the PA-TM-RING E3 ligase itself, either by homodimerization or heterodimerization between two ligases. Using this approach would allow strategic modulation of transmembrane E3 ligases and consequently protein homeostasis of their natural targets, a process we termed Fratricide REULRs (Figure 5B). Indeed, treatment of cells with RNF128, RNF130, RNF167, RNF43 and ZNRF3 Fratricide REULR molecules resulted in effective loss of cell surface ligase levels in HEK293T cells (Figure 5C-D, Figure 5-figure supplement 1A).

Furthermore, to demonstrate the modular nature and flexibility of the nanobody based REULR design, we engineered a RNF43-ZNRF3 heterobifunctional REULR (Figure 5B, Figure 5-figure supplement 1B) that would allow the elimination of two ligases using one Fratricide REULR molecule. Treatment of HEK293T cells overexpressing MYC-tagged RNF43 and HA-tagged ZNRF3 with a RNF43-ZNRF3 heterobifunctional Fratricide REULR (Figure 5D; right bar graph) resulted in significant reduction of RNF43 and near compete loss of ZNRF3 levels comparable to RNF43 and ZNRF3 Fratricide REULRs (Figure 5D; left and middle bar graph).

To demonstrate the ease of formatting using PA-TM-RING E3 binding VHHs, we extended the previous design into a linear, hetero-trimeric array of VHHs targeting RNF128, RNF43 and ZNRF3 with one Fratricide REULR molecule (Figure 5E, Figure 5-figure supplement 1C). We co-expressed HA-tagged RNF128, MYC-tagged ZNRF3 and FLAG-tagged RNF43 in HEK293T cells and treated cells with RNF128 (only targets RNF128) or ZNRF3-REULR (only targets ZNRF3), heterobifunctional RNF43-ZNRF3 REULR (targets RNF43 and ZNRF3) or a hetero-trimeric RNF43-RNF128-ZNRF3 Fratricide REULR that targets all three PA-TM-RING E3 ligases for degradation (Figure 5F). A RNF43-RNF128-ZNRF3 targeting VHH array was able to efficiently eliminate all three E3 Ligases from the cell surface and further shows the robustness and advantage of a “mix and match” VHH based targeting approach.

The WNT signaling pathway is instrumental for embryonic development, stem cell differentiation and regeneration of injured tissues and modulation of WNT signaling presents an untapped potential in regenerative medicine (Barker and Clevers, 2006; Logan and Nusse, 2004; MacDonald et al., 2009; Niehrs, 2012). RNF43 and ZNRF3 are two pivotal PA-TM-RING E3 ligases known to negatively regulate the WNT signaling pathway by targeting Wnt receptors FZD and promote receptor degradation via the UPS (Figure 6A) (Clevers and Nusse, 2012; Zebisch and Jones, 2015). With well-established Fratricide REULRs in hand, we explored whether RNF43 and ZNRF3 based Fratricide REULR molecules have the potential to modulate FZD receptor cell surface levels and potentiate downstream WNT signaling events. We first treated HEK293T cells with RNF43 or ZNRF3 Fratricide REULR molecules and monitored FZD cell surface levels using a previously developed pan-FZD (DRPB_Fz7/8) as a staining reagent due to its high affinity and broad binding spectrum for FZD receptors: FZD1, 2, 5, 7 and 8 (Dang et al., 2019). We indeed observed significant increase in the accumulate of FZD levels after RNF43 or ZNRF3 Fratricide REULR treatment compared to PBS or monomeric RNF43 or ZNRF nanobodies (Figure 6B). To examine whether these results can be translated into a functional assay and elicit Fratricide REULR specific activation of canonical WNT signaling, we performed a series of reporter assays using HEK 293STF (SuperTopFlash) cells. In agreement with increased FZD levels upon treatment with RNF43 or ZNRF3 Fratricide REULRs we similarly observed robust induction of WNT signaling and increased signaling activity using a heterospecific RNF43-ZNRF3 Fratricide REULR, compared to treatment with WNT or monomeric PA-TM-RING nanobodies alone (Figure 6C).

**Figure 6.**
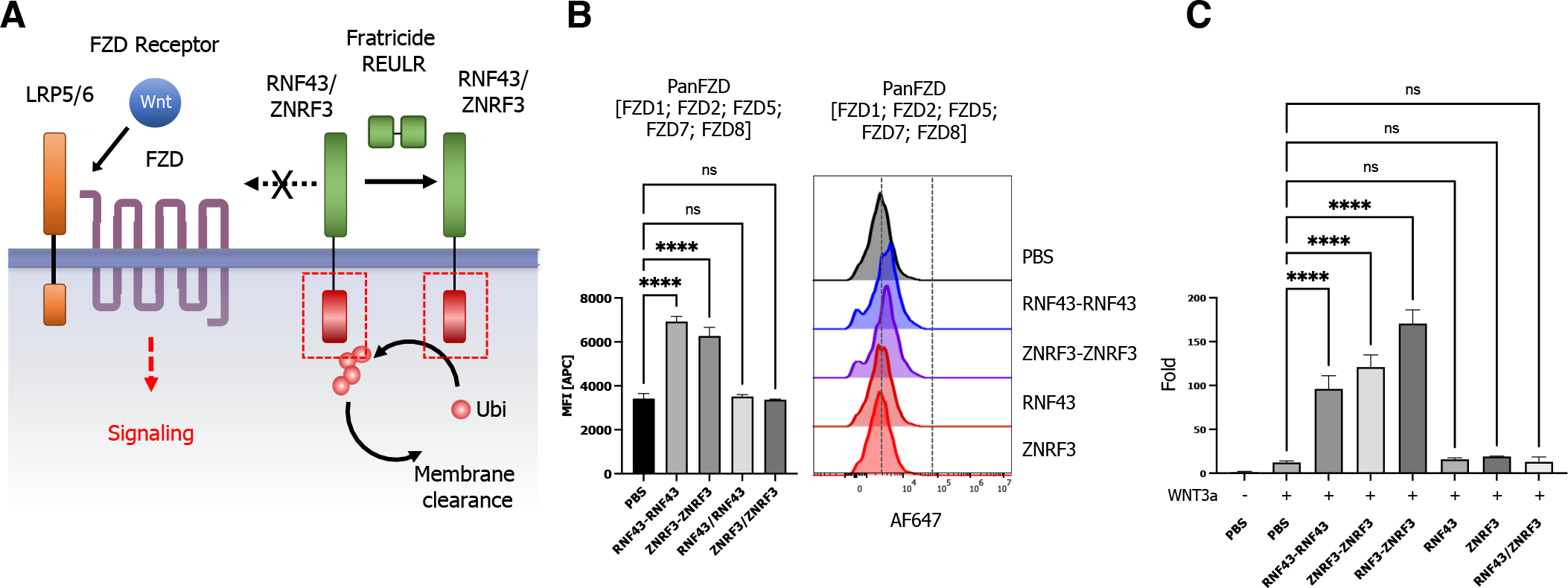
Fratricide REULR and WNT Signaling potentiation. (A) Schematic representation of a RNF43 or ZNRF3 based Fratricide REULR in the context of canonical WNT signaling pathway. (B) HEK293STF cells were seeded at 10k/well and subsequently treated with RNF43 or ZNRF3 Fratricide REULR for 24h. FZD cell surface levels were measured by incubating cells with a biotinylated pan-FZD Darpin (DRPB_Fz7/8), recovered by SA647 and analyzed by flow cytometry. Data are mean ± s.d. (n = 3 replicates). (C) HEK293STF cells were seeded at 5k/well and treated with RNF43, ZNRF3 or RNF43-ZNRF3 Fratricide REULR molecules after 24h in the presence of 10% conditioned WNT3a media (monomeric binding moieties or PBS were used as negative controls). After 36h, activation of the β-catenin dependent STF reporter by Fratricide REULRs was measured. Data are mean ± s.d. (n = 3 replicates).

## Discussion

In summary, we implemented a modular, “mix and match” human and mouse cross-reactive nanobody based targeted protein degradation platform termed REULR (Receptor Elimination by E3 Ubiquitin Ligase Recruitment) by retasking 5 PA-TM-RING E3 Ligases (RNF128, RNF130, RNF167, RNF43 and ZNRF3) to modulate cell surface receptors by induced proximity allowing selective, tissue specific application. REULR based bispecific molecules can be broadly applied to modulate cell surface levels of a variety of therapeutically relevant transmembrane receptors using different binding moieties (Figure 2-4). Furthermore, we present a strategic approach to tune transmembrane E3 Ligases itself by using homo- and heterobispecific Fratricide REULRs and consequently modulate signaling events of natural target receptors (Figure 5, 6).

While similar approaches (AbTACs, PROTABs) have been reported during the course of this study to degrade PD-L1 or IGF1R, they are mainly limited by using WNT responsive E3 ligase RNF43 and ZNRF3 (Cotton et al., 2021; Marei et al., 2022) and rely on human IgG antibody scaffolds to generate heterobifunctional targeting molecules. Antibody-derived biologics are generally more constrained by their inherent structural properties including large size (150 kDa), formatting and modularity that might limit the applicability for tumor therapy. By contrast, REULR molecules take advantage of the superior pharmacokinetic properties of nanobodies (VHH) allowing for a versatile and modular design with ease of formatting into homo- or heterobifunctional dimer or arrayed multimers to target one or multiple cell surface proteins (Figure 2-5).

Interestingly, we observed that the affinity and expression levels of the PA-TM-RING targeting nanobody did not directly correlate with levels of cell surface clearance, suggesting that the PA-TM-RING E3 Ligases operate with a wide spectrum of cytosolic catalytic RING E3 activity rather than abundance. Morevover, REULR processivity may be further influenced by orientation and geometry of the ternary Receptor-REULR-Ligase complex.

In addition to the therapeutic potential of REULR molecules, the monomeric binding modules (nanobodies) itself present invaluable tools to validate natural targets and to gain a deeper understanding into the fundamental biological function of transmembrane E3 ligases and their cellular pathways in drug discovery and the context of cancer biology.

Collectively, we believe that our “mix and match” nanobody based REULR protein degradation strategy holds tremendous promise for a large variety of targets and serve as powerful research tools with the potential to novel therapeutic applications that can be easily customized by virtue of it’s modularity, human and mouse cross reactivity and tissue specificity.

## MATERIALS AND METHODS

### Curation of the human ubiquitin cell surface receptor proteome

A raw list of reported ubiquitin sites was obtained from PhosphoSitePlus (PSP; https://www.phosphosite.org) and matched to a curated list of the human membrane proteome (Siepe et al., 2022) to generate a master list of cell surface receptors with reported ubiquitination sites.

### Database integration

Pairwise protein sequence alignments were performed using Smith-Waterman algorithm to calculate alignments between human and mouse amino acid sequences obtained from UniProt (https://www.uniprot.org/). Phylogenetic homology analysis (PHA) was performed to generate phylogenetic trees from multiple sequence alignments (MSA) of amino acid sequences of secreted or ECD sequences of transmembrane cell surface receptors (https://www.uniprot.org/). Briefly, MSA was performed using ClustalOmega (https://www.ebi.ac.uk/Tools/msa/clustalo/) and alignments results were submitted to calculate phylogenetic tree parameters (https://www.ebi.ac.uk/Tools/phylogeny/simple phylogeny/) which were visualized by Interactive Tree of Life (iTOL; https://itol.embl.de/) (Letunic and Bork, 2021). Tissue expression datasets, normal tissue and TCGA (The Cancer Genome Atlas) datasets, were downloaded from The Human Protein Atlas (https://www.proteinatlas.org; v21.1). TCGA cancer tissue RNA-seq data was obtained from 17 cancer types representing 21 cancer subtypes and were processed as median FPKM (number Fragments Per Kilobase of exon per Million reads) and visualized as a hierarchical clustering heatmap using JMP Pro (v16). Unsupervised hierarchical clustering of normalized mRNA gene expression by tissue was performed with Ward linkage and correlation distance were plotted as heatmaps using JMP Pro (v16).

### Cell lines

Suspension cells were grown in plain bottom, vented flasks (Thermo), adherent cells were grown in T25 or T75 flasks (ThermoFisher). Cells were maintained at 37 °C and 5% CO2. HEK293T (CRL-3216; ATCC) and LentiX cells were maintained in DMEM supplemented with 10% FBS, 1% GlutaMax and 1% penicillin/streptomycin. HEK293F (R79007; ThermoFisher) were grown in FreeStyle media (12338018; ThermoFisher). Expi293F (A14528; ThermoFisher) cells were grown in Expi293™ Expression Medium (ThermoFisher). Cell lines tested negative for mycoplasma (MycoAlert Mycoplasma Detection kit, Lonza).

### Facs staining

Cells were stained with the indicated antibodies at 1:100 dilution or tetramer at the indicated concentration for 30 min on ice in MACS staining buffer (Miltenyi). After incubation with fluorescent antibodies, cells were washed with MACS buffer and analyzed via flow cytometry on a Cytoflex (Beckman Coulter) instrument. Surface expression was quantified by FACS using the CytoFLEX equipped with a high-throughput sampler. Live cells were identified after gating on the basis of forward scatter (FSC) and side scatter (SSC) and propidium iodide (PI)-negative staining. Data were analyzed using FlowJo 10.8.1 (BD). All assays were performed using independent biological replicates. The number of replicates (n) is indicated in the figure legends. Mean fluorescence intensity (MFI) was determined in FlowJo 10.8.1.

### Antibodies

Primary antibodies used in this study include anti-DYKDDDDK Tag (CST, D6W5B, # 15009), anti-HA Tag (CST, 6E2, #3444), anti-MYC (CST, 9B11, #2279). Antibodies were used at 1:100 dilution in MACS staining buffer (Miltenyi).

### Production of purified proteins

Proteins were produced in Expi293F cells using transfection conditions following the manufacturer’s protocol. After harvesting of cell media, 1 M Tris, pH 8.0 was added to a final concentration of 20 mM. Ni-NTA Agarose (Qiagen) was added to ~5% media volume. 1 x sterile PBS, pH 7.2 (Gibco) was added to ~3X media volume. The mixture was stirred overnight at 4° C. Ni-NTA agarose beads were collected in a Buchner funnel and washed with ~300 ml protein wash buffer (20 mM HEPES, pH 7.2, 150 mM NaCl, 20 mM imidazole). Beads were transferred to an Econo-Pak Chromatography column (Bio-Rad) and protein was eluted in 15 ml of elution buffer (20 mM HEPES, pH 7.2, 150 mM NaCl, 200 mM imidazole). The DNA encoding for pan-FZD (DRPB_Fz7/8) was cloned into pET-28 with a C-terminal AVI-6xHIS tag and transformed into Rosetta DE3 competent cells. The cells were grown at 37°C in 2YT media supplemented with Kanamycin (40 μg/mL) until the culture reached log-phase growth. IPTG was added to the culture to induce protein expression at a final concentration of 1 mM. The culture was shaken at 37°C for 3 hours and protein was harvested from the cells by sonication. Pan-Fzd protein was purified using Ni-NTA Agarose (Qiagen), followed by biotinylation and size-exclusion chromatography with a Superdex S75 column (GE Healthcare). In general, proteins were concentrated using Amicon Ultracel filters (Millipore) and absorbance at 280 nm was measured using a Nanodrop 2000 spectrophotometer (Thermo Fisher Scientific) to determine protein concentration.

### Biotinylation and FPLC Purification

Where indicated, proteins were biotinylated as described previously (Özkan et al., 2013). Briefly, up to 10 mg of protein was incubated at 4° overnight in 2X Biomix A (0.5 M bicine buffer), 2X Biomix B (100 mM ATP, 100 mM MgOAc, 500 μM d-biotin), Bio200 (500 uM d-biotin) to a final concentration of 20 uM, and 60-80 units BirA ligase in a final volume of 1 ml. All proteins were further purified by size-exclusion chromatography using an S75 or a S200 Increase column (GE Healthcare), depending on protein size, on an ÄKTA Pure FPLC (GE Healthcare), FPLC traces for purified proteins used for SPR can be found in Figure 2-figure supplement 2 and S8=Figure 2-figure supplement 3.

### Nanobody selection

Nanobody selection was performed as previously described with minor alterations. Briefly, the synthetic yeast library was expanded overnight in -Trp media with glucose at 30C and induced at 10x the theoretical diversity by suspension in -Trp media with galactose, grown at 20C for 24h. Surface display was assessed by flow cytometry after staining with an anti-HA antibody. Round 1 and 2 were first negatively selected on magnetic streptavidin beads and then positively selected on magnetic streptavidin beads loaded with biotinylated target protein. Subsequent rounds were carried out with target protein tetramerized by streptavidin and bound to anti-fluorophore magnetic beads followed by decreasing monomer protein concentration and by flow cytometry. Single clones from the final round were sorted into 96 well plates, induced for 24h at 20C and grown in deep well blocks. The top 20 clones were sequenced and unique clones were then expressed in Expi293F cells and assayed for binding to the target protein by SPR.

### Cell-Surface binding assay with streptavidin tetramerized proteins

To examine PPIs at the cell surface, we performed cell-surface protein binding assays using HEK293F, HEK293T or PBMC cells. HEK293F or HEK293T cells were transfected using Fugene6 according to manufacturer’s protocol (Promega) with expression plasmids encoding full-length proteins containing an N-terminal tag (FLAG, HA or MYC). Two days following transfection, cultures were harvested, cells were spun down for 4 min at 1600 rpm (~400x g), washed twice with cold MACS buffer (Miltenyi) and resuspended to a final density of ~3 x 10^6^ cells/ml. To generate tetramerized proteins to test for binding to cells expressing full-length proteins, FPLC-purified biotinylated proteins (see above) were incubated with streptavidin tetramers conjugated to Alexa647 Fluor (SA-647) (Thermo Fisher) at a 4:1 molar ratio on ice for at least 15 minutes. To assess cell-surface expression of full-length cell surface receptors, anti-FLAG (CST), anti-HA (CST) or anti-MYC antibody (CST) staining of cells was also performed in parallel at 1:200 with either Alexa647 or Alexa488 conjugated antibodies, as indicated. Approximately 150,000 cells were incubated with Protein:SA-647 complexes or antibody in a final volume of 100 ul in 96-well round-bottom plates (Corning) for 1 hour at 4° C protected from light. Following incubation, cells were washed two times with 200 ul cold MACS buffer and resuspended in 200 ul cold MACS buffer with 1:3000 propidium iodide (Thermo Fisher Scientific). Immunofluorescence staining was analyzed using a Cytoflex (Beckman Coulter), and data were collected for 20,000 cells. Data were analyzed using FlowJo v10.4.2 software. All data report mean fluorescence intensity (MFI). Concentration-dependent binding of Protein:SA-647 to full-length receptor-expressing, but not mock control cells, was deemed indicative of cell-surface binding.

### SPR Experiments

SPR experiments were performed using a Biacore T100 instrument (GE Healthcare). FPLC-purified biotinylated proteins (ligands) in HBS-P+ Buffer (GE Healthcare) were captured on a Streptavidin (SA) Series S Sensor Chip (GE Healthcare). Chip capture was performed in HBS-P+ buffer (GE Healthcare) to aim for ~100-200 ligand response units (RU). Flow cell 1 was left empty to use this flow cell as a reference flow-cell for on-line subtraction of bulk solution refractive index and for evaluation of non-specific binding of analyte to the chip surface using Biacore T100 Control Software (version 3.2) (GE Healthcare). FPLC-purified non-biotinylated protein was used as analyte. Analytes were run in HBS-P+ buffer using 2-fold increasing protein concentrations to generate a series of sensorgrams. Binding parameters were either determined based on a 1:1 Langmuir model or at equilibrium using the accompanying Biacore T100 evaluation software.

### STF luciferase reporter assays

HEK293STF cells were seeded for each condition in 96-well plates, and stimulated with Fratricide REULRs, WNT (WNT3a conditioned media; ATCC), control proteins, or PBS for 36h. After washing cells with 1xPBS, cells in each well were lysed in 30 μl 1× passive lysis buffer (Promega). 10 μl per well of lysate was assayed using the Dual Luciferase Assay kit (Promega).

### Statistics

All figures are representative at least n=3 (in vitro) experiments unless otherwise noted. Statistical significance was assayed by grouped, one-way ANOVA using GraphPad Prism 9.4.1. In all figures *P < 0.05; **P < 0.01; ***P < 0.001; ****P < 0.0001; NS: not significant. Data are represented as mean ± s.d., unless otherwise stated.

## Data Availability

Data that support the findings of this study are available from the corresponding author upon request.

## Author Contributions

Conceptualization, D.H.S. and K.C.G.; Methodology, D.H.S. and K.C.G.; Nanobody library screening and protein expression for in vitro studies, D.H.S. and L.K.P.; Analysis, D.H.S.; Investigation, D.H.S. and K.C.G.; Writing – Original Draft, D.H.S; Writing – Review & Editing, D.H.S. and K.C.G.; Visualization, D.H.S.; Supervision, K.C.G.; Funding Acquisition, K.C.G.

## Acknowledgments

The authors are funded by the Howard Hughes Medical Institute, 2R01CA177684 and Emerson Collective, (K.C.G.).

---

**Figure 1-figure supplement 1.**
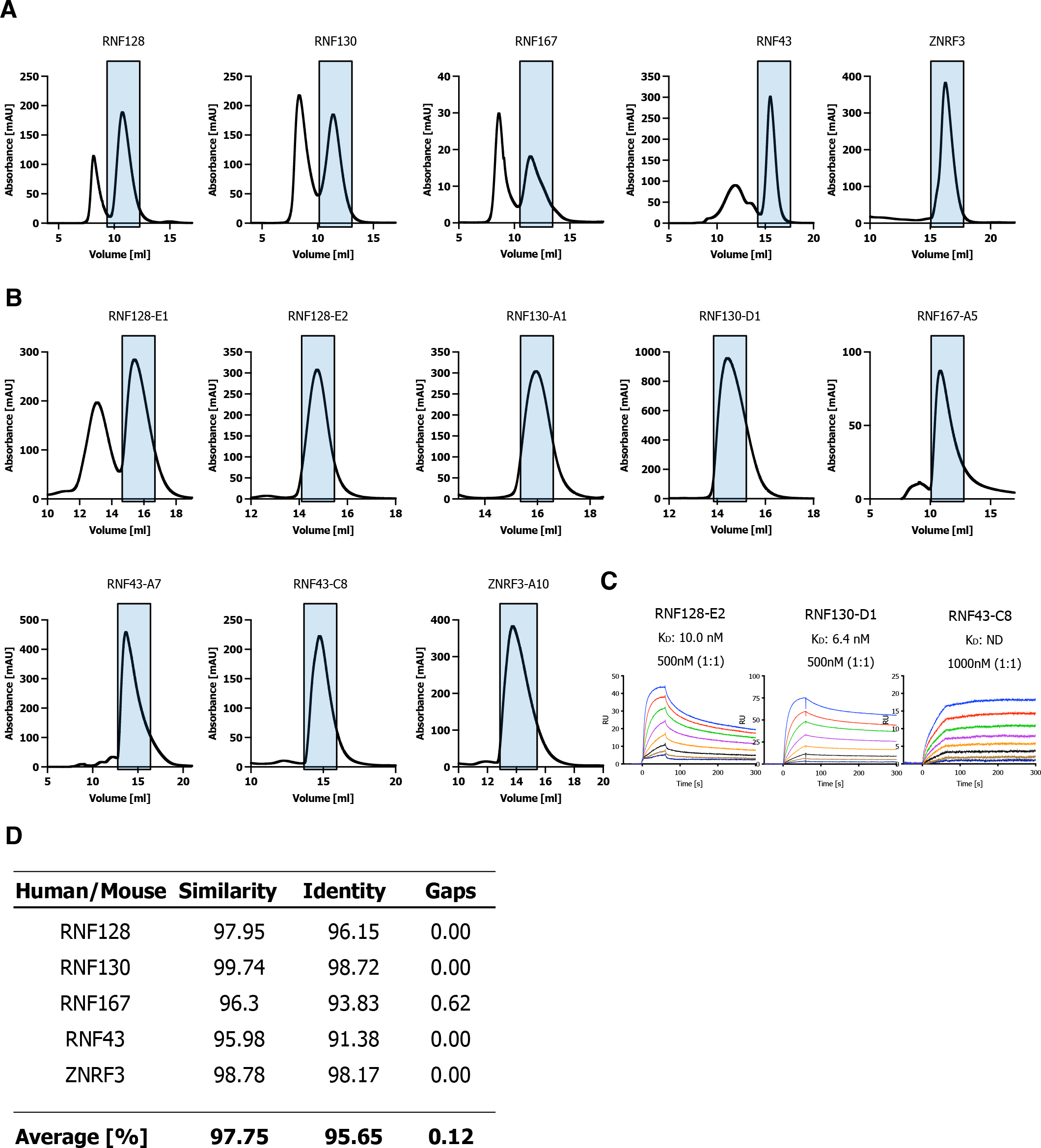
Size-Exclusion Chromatography. (A, B) Size-Exclusion Chromatography traces of RNF128, RNF130, RNF167, RNF43 and ZNRF3 PA-TM-RING E3 Ligase ECDs used for screening the synthetic nanobody library and SPR validation of PA-TM-RING E3 Ligase VHHs. (C) SPR sensograms and binding affinities of PA-TM-RING ligase selected nanobodies (analytes) for human RNF128, RNF130 and RNF43 ECDs (ligands). (D) Pairwise protein sequence alignments of PA-TM-RING E3 Ligases were performed using Smith-Waterman algorithm to calculate alignments between human and mouse amino acid sequences.

**Figure 1-figure supplement 2.**
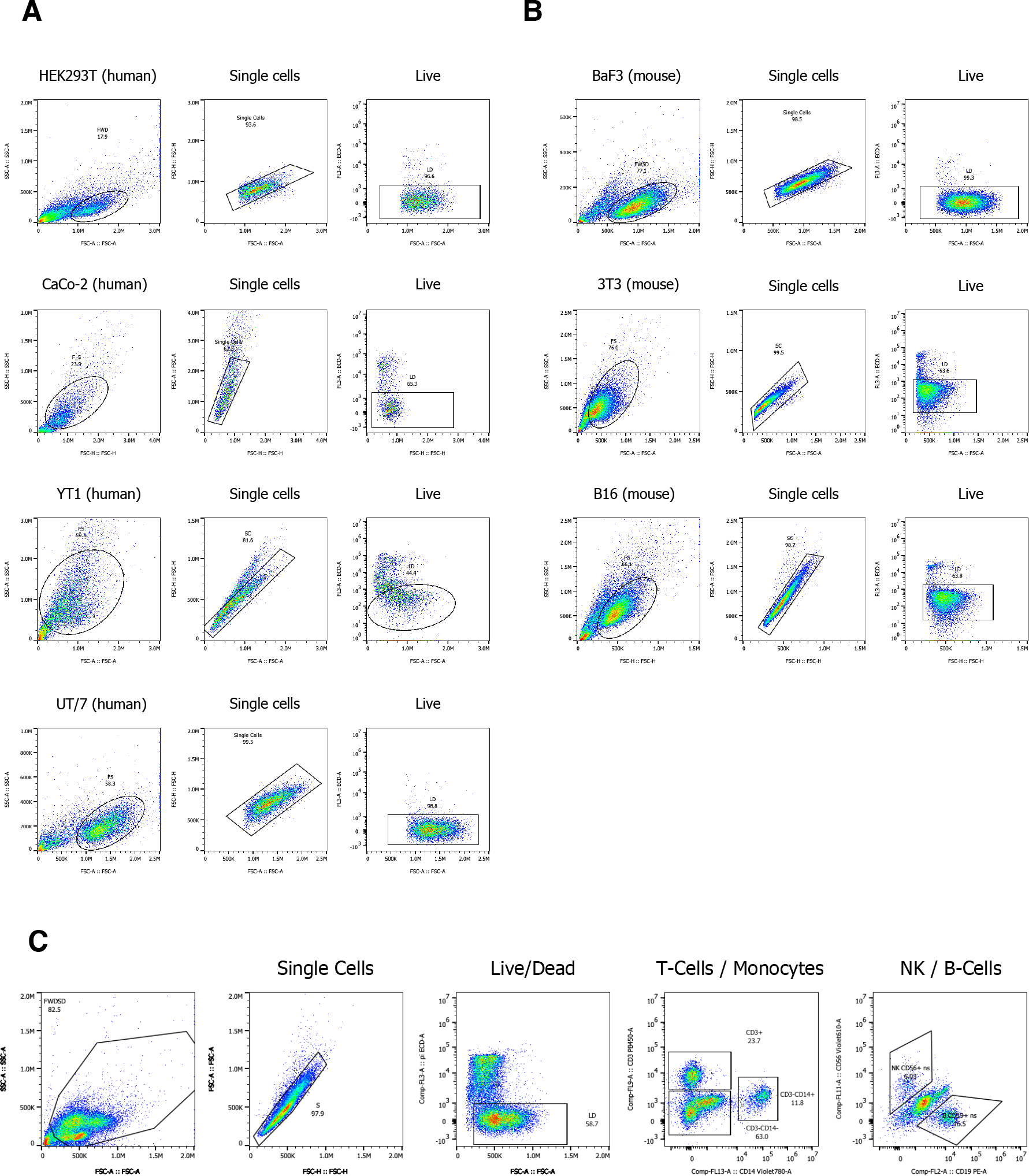
Cell Surface Staining. (A) Gating strategy of human HEK293T, CaCo-2, YT1, UT/7 cell lines. (B) Gating strategy of mouse BaF3, 3T3, B16 cell lines. (C) Gating strategy of Primary Peripheral Blood Mononuclear Cells (PBMCs).

**Figure 2-figure supplement 1.**
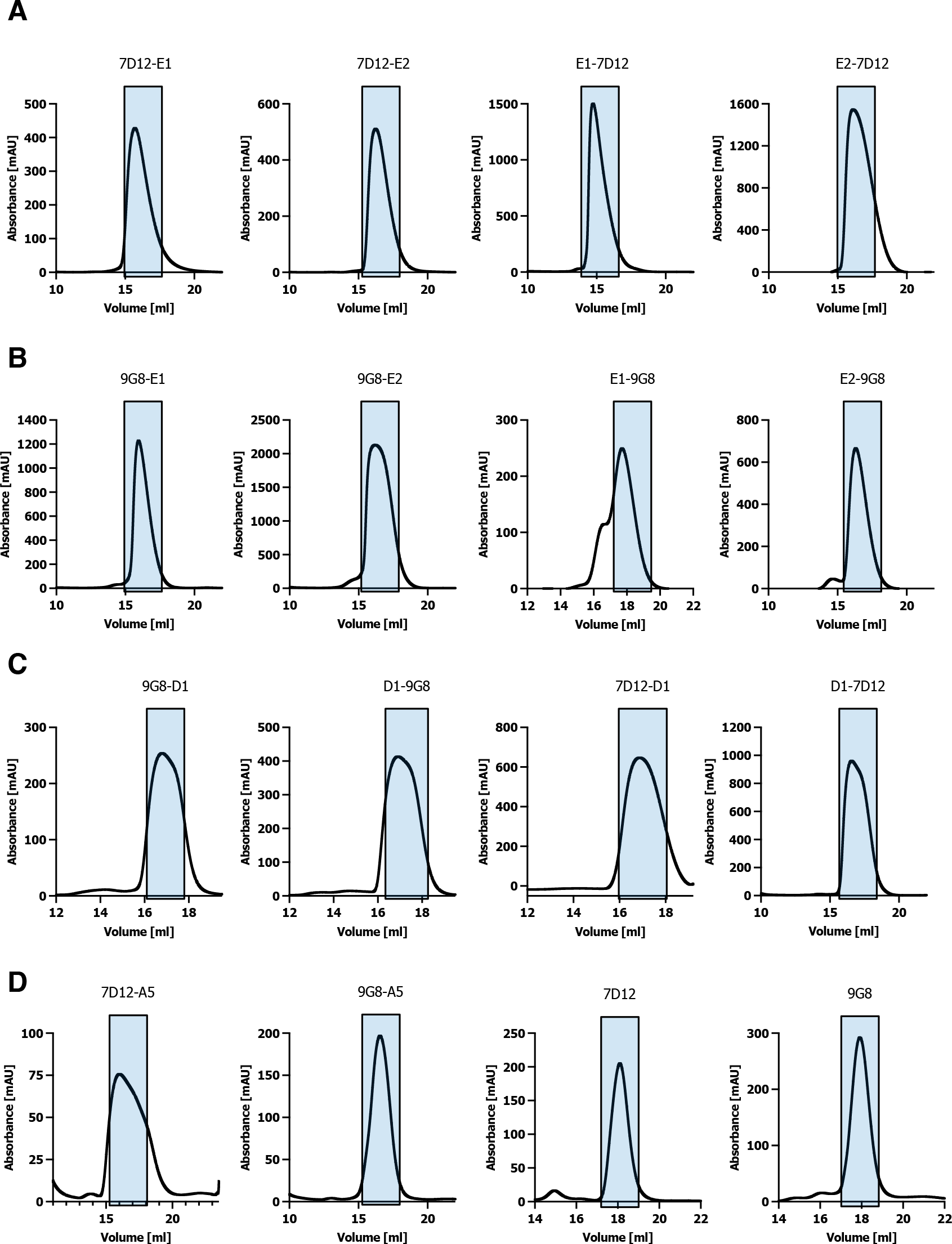
Size-Exclusion Chromatography. (A-D) Size-Exclusion Chromatography traces of proteins used for EGFR-REULR experiments.

**Figure 3-figure supplement 1.**
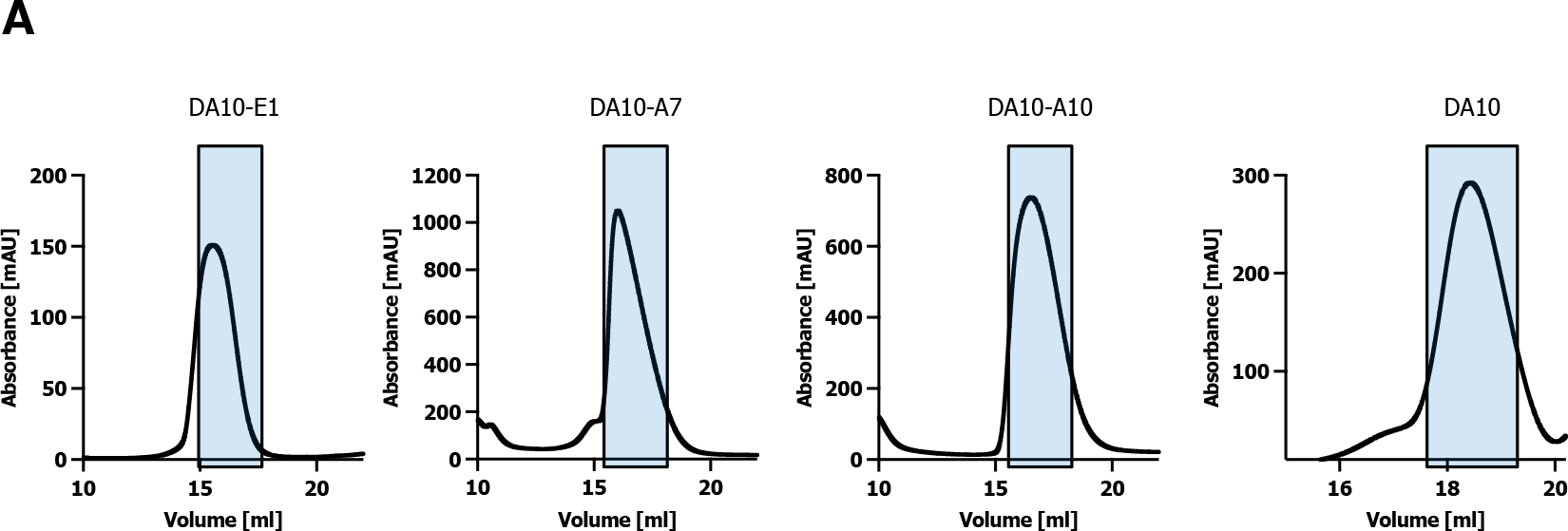
Size-Exclusion Chromatography. (A) Size-Exclusion Chromatography traces of proteins used for EPOR-REULR experiments.

**Figure 4-figure supplement 1.**
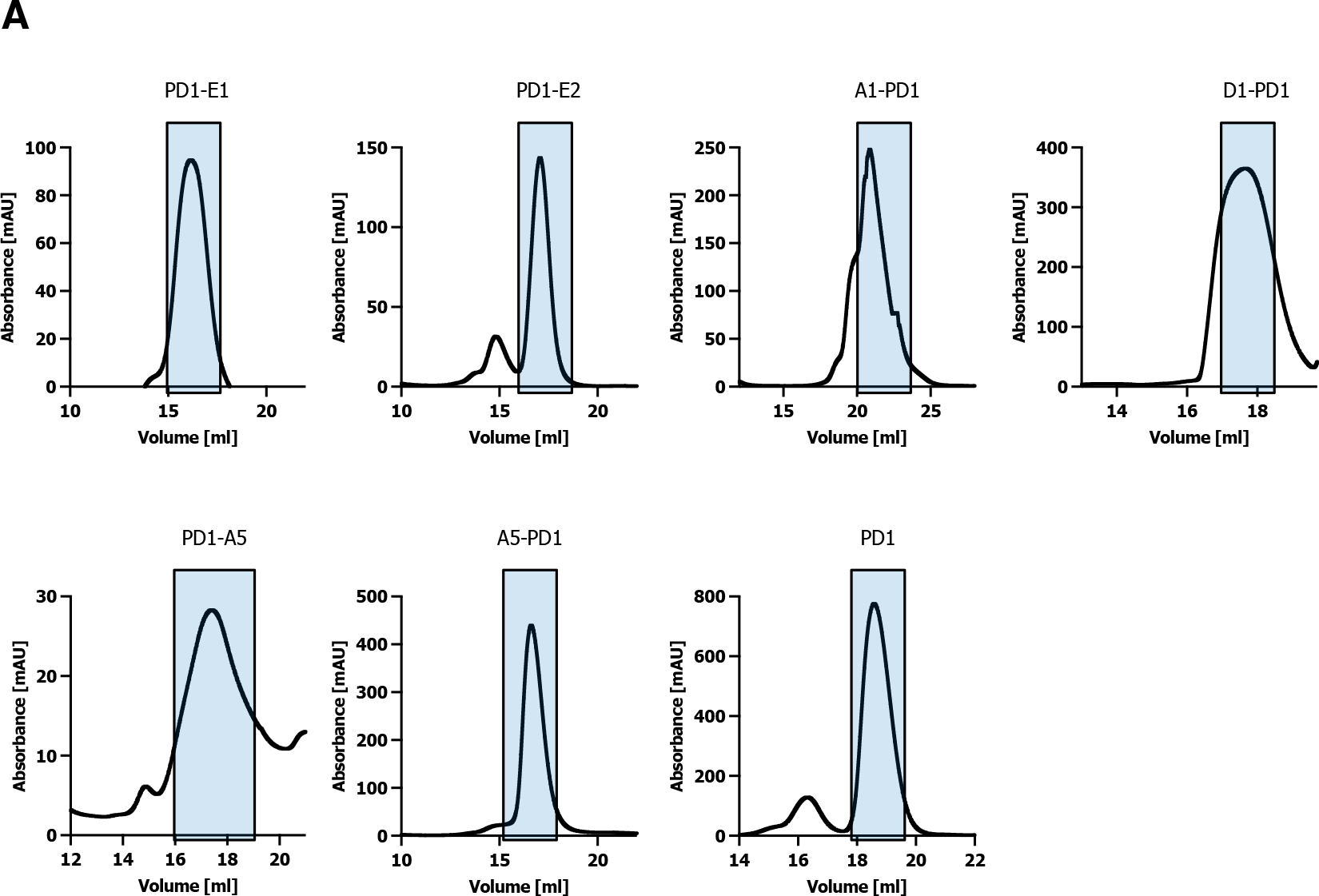
Size-Exclusion Chromatography. (A) Size-Exclusion Chromatography traces of proteins used for PD1-REULR experiments.

**Figure 5-figure supplement 1.**
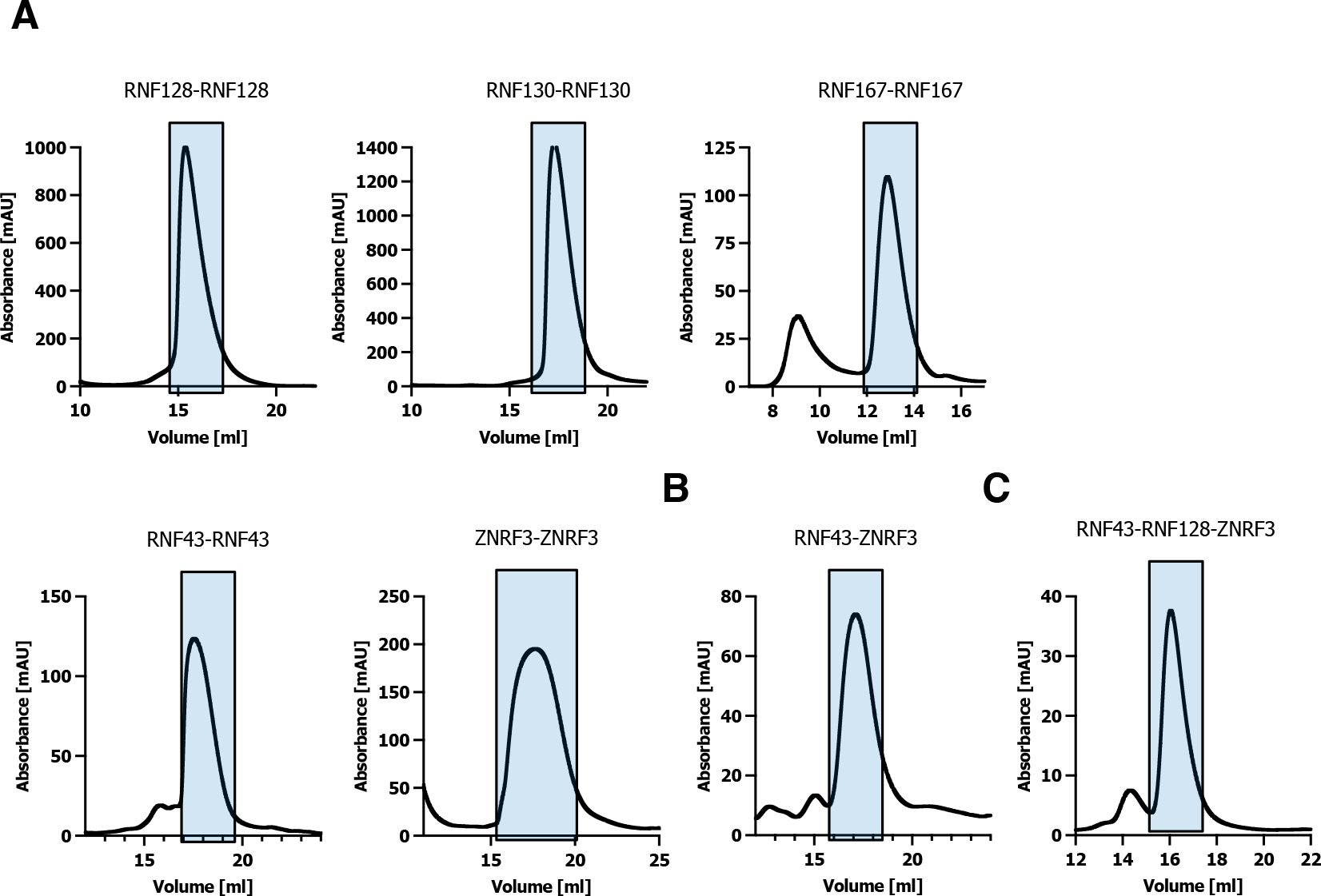
Size-Exclusion Chromatography. (A-C) Size-Exclusion Chromatography traces of proteins used for Fratricide REULR experiments.

